# Convergent evolution of complex adaptive traits enabled human life at high altitudes

**DOI:** 10.1101/2023.10.24.563738

**Authors:** Giulia Ferraretti, Aina Rill, Paolo Abondio, Kyra Smith, Claudia Ojeda-Granados, Sara De Fanti, Massimo Izzi, Phurba T. Sherpa, Paolo Cocco, Massimiliano Tiriticco, Marco di Marcello, Agnese Dezi, Guido Alberto Gnecchi-Ruscone, Luca Natali, Angela Corcelli, Giorgio Marinelli, Paolo Garagnani, Davide Peluzzi, Donata Luiselli, Davide Pettener, Stefania Sarno, Marco Sazzini

**Author notes:** These authors equally contributed (Giulia Ferraretti, Aina Rill, Paolo Abondio). These authors equally contributed (Stefania Sarno, Marco Sazzini).

## Abstract

Convergent adaptations represent paradigmatic examples of the capacity of natural selection to influence organisms’ biology. However, the possibility to investigate genetic determinants underpinning convergent complex adaptive traits has been offered only recently by methods for inferring polygenic adaptations from genomic data. Relying on this approach, we demonstrate how high-altitude Andean human groups experienced pervasive selective events at angiogenic pathways, which resemble those previously attested for Himalayan populations despite partial convergence at the single-gene level was observed. This provides unprecedented evidence for the drivers of convergent evolution of enhanced blood perfusion in populations exposed to hypobaric hypoxia for thousands of years.

## Introduction

Convergent evolution refers to an evolutionary scenario in which different species or populations independently develop the same (or similar) biological trait instead of inheriting it from a common ancestor. In particular, convergent adaptation occurs when distantly related species/populations, which live in distinct geographical areas or epochs, occupy comparable ecological settings and are subjected to the same selective pressures, thus being similarly targeted by natural selection (Stern, 2013). Several cases of convergent evolution have been described so far in a number of animal and plant taxa (Scott & Spielman, 2006; Stern, 2013; Chemes et al., 2015; Wen et al., 2020), with fewer examples (e.g., skin pigmentation, malaria resistance, lactose tolerance, and metabolic adaptations to cold climates) being instead reported in the human species when comparing adaptive traits among populations of different ancestry (Kwiatkowski, 2005; Edwards et al., 2010; Balentine & Bolnick, 2022). Nevertheless, experimental approaches to accurately characterize genomic variability within and between species/populations, as well as inferential statistical methods suitable to investigate how far phenotypic convergence is achieved by means of genetic convergence, have been only recently conceived. This is especially the case of complex (i.e., polygenic) adaptive traits, whose evolution is regulated by changes at several genes that are functionally related and contribute to the modulation of the same biological function (Pritchard & Di Rienzo, 2010). Development of methods able to test the occurrence of selective events under a realistic approximation of a polygenic adaptation model has thus opened new possibilities to assess whether natural selection acted on the same genetic variants or genes or functional pathways in species/populations showing shared or similar adaptive traits (Gouy et al., 2017).

To this end, human adaptation to the high-altitude environment represents a valuable case study because the main selective pressure acting on populations dwelling at altitude (i.e., hypobaric hypoxia) cannot be mitigated by cultural adaptations and thus acts with the same intensity on human groups living at comparable altitudes irrespectively to their ancestry, geographical locations, and socio-cultural contexts. We can thus hypothesize that Himalayan and Andean high-altitude populations, which represent the main human groups whose ancestors have had to cope with such a selective pressure, experienced convergent evolution at least to some extent. Despite this assumption, limited overlapping of potential adaptive loci (e.g., entailing variants at the *EGLN1, EDNRA*, and *NOS2* genes) has been described so far between them (Ferraretti et al., n.d.; Witt & Huerta-Sánchez, 2019). This may be partially due to the limitations imposed by studies that to date focused mainly on selective sweeps. Indeed, these adaptive mechanisms are mediated by the action of natural selection on single/few genetic variants, which implies that these loci can exert a large effect on a given biological trait (Pritchard & Di Rienzo, 2010). That being so, selective sweeps have low probability to occur in human populations, who have long maintained low effective population sizes and have accumulated low genetic variability, and thus need several thousands of years to evolve (Hernandez et al., 2011). Coupled with the considerably more recent human colonization of the high-altitude ecological niche in the Andes than in the Himalayas, as supported by estimates at around 9 kya for the divergence between Andean and lowland South American populations (Lindo et al., 2018) and by archeological evidence for occupation of the Tibetan Plateau since Paleolithic times (Zhang et al., 2021), this suggests selective sweeps might be not the predominant drivers of adaptation of Andean people to such a challenging environment.

Consistent with this scenario, Andeans indeed evolved less effective genetically regulated physiological adjustments, especially to counteract long-term detrimental effects of hypobaric hypoxia, than populations of Tibetan/Sherpa ancestry. When compared with acclimatized Native American lowlanders, they exhibit traits contributing to enhanced efficiency of respiratory functions and oxygen utilization, such as larger lung volumes and chest dimensions, narrower alveolar to arterial oxygen gradients, less hypoxia-induced pulmonary vasoconstriction, greater uterine artery blood flow during pregnancy, less altitude-related decrement in maximal exercise oxygen consumption and increased cardiac oxygen utilization (Beall, 2007; Julian & Moore, 2019). However, when genetic determinants of these physiological characteristics have been searched for, most of the inferred selective sweeps failed to be replicated by multiple studies, suggesting that their relationships with the observed adaptive traits are far from being fully elucidated (Bigham, 2016; Crawford et al., 2017; Lindo et al., 2018; Jacovas et al., 2018; Julian & Moore, 2019).

## Results and Discussion

To test whether polygenic adaptations played a role in shaping the Andean adaptive phenotype, and to extend to complex traits the investigation of the genetic bases of potential convergent evolution between Andean and Himalayan populations, we assembled a dataset representative of genomic variation at several high-altitude ethnic groups from the Andean region (Supplementary Table 1). After stringent quality control (QC) filtering (see Methods), 7,966,198 single nucleotide variants (SNVs) from 24 Bolivian Aymara whole genome sequences (WGS) from Click or tap here to enter text. were used to perform multiple selection scans, whose results informed gene-network analyses aimed at identifying sets of functionally related loci enriched for signatures of natural selection. Imputed genotype data for 8,024,216 SNVs from 130 individuals belonging to additional Bolivian Aymara groups, as well as from Aymara, Quechua, and Uros high-altitude Peruvian populations (Reich et al., 2012; Gnecchi-Ruscone et al., 2019) instead served as validation datasets.

### Identification of Native American Low-altitude Control Populations

Setting the considered Bolivian Aymara WGS into the genomic landscape of indigenous groups from Central and South America enabled us to confirm them as well representative of the genetic variation observable across Andean high-altitude populations (Fig. 1a, Supplementary Results). Comparable results were obtained also when considering imputed data for Andeans, thus providing evidence for consistency between non-imputed and imputed datasets (Supplementary Fig. 1). Such a consistency was further attested by the high and significant correlation observed between pre- and post-imputation identity-by-state (IBS) matrices of individual pairwise differences in the genome-wide proportion of shared alleles (Mantel correlation = 0.95, P < 10^−4^).

**Figure 1.**
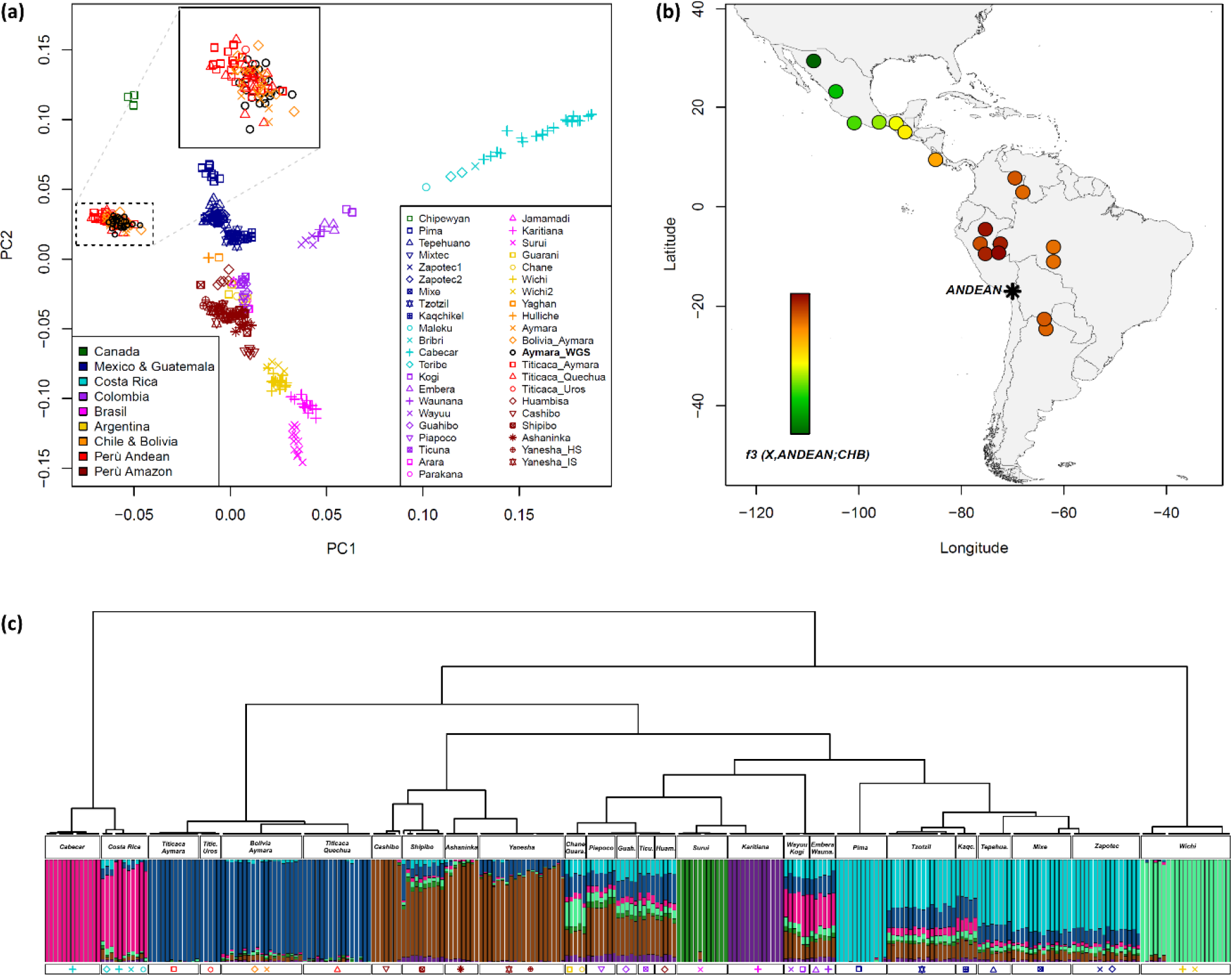
Population structure analyses performed to investigate genetic relationships between high-altitude Andean groups and low-altitude Central and South American populations. **(a)** PCA representing PC1 versus PC2 computed for the 43 un-admixed Native American groups reported in the legend at the bottom right of the plot. Individuals are color-coded according to their country of origin as reported in the bottom-left legend. The box at the top of the plot details the position of Bolivian Aymara WGS in the considered genetic space with respect to the other high-altitude Andean populations. **(b)** Heat map of values for the outgroup-f3 statistic representing the estimated amount of shared genetic drift between the Andean population cluster (marked by a star-like symbol) and each of the considered populations from Central and Southern America (indicated in their approximate geographic locations) A gradient ranging from green to red is specifically used to indicate lower-to-higher levels of genetic affinity as reported by the corresponding color scale. **(c)** Fine-scale patterns of genetic clustering and proportions of ancestry components inferred at the individual level. The fineSTRUCTURE hierarchical clustering dendrogram obtained for Andean, Central and South American individuals included in the assembled dataset is reported, along with ancestry components inferred for each subject with ADMIXTURE analysis at K = 8. For each genetic cluster pinpointed by the fineSTRUCTURE analysis, the corresponding individuals and their admixture proportions are specifically detailed, along with the symbols used in the PCA plot for the populations they belong to.

The outgroup-f3 statistic was then computed on the assembled reference dataset to select the most suitable low-altitude control populations of South American ancestry to be contrasted with Andean groups in terms of evolved biological adaptations (see Materials and Methods for a detailed description of the adopted selection criteria). According to this approach, while each Andean population was confirmed to share the greatest genetic similarity with the other Andean groups, Amazonian populations from Peru appeared as the next most affine ones to Andeans (Fig. 1b, Supplementary Fig. 2 and Supplementary Results). Among these putative control populations, 24 Cashibo, Shipibo, and Ashaninka subjects were specifically selected to constitute the low-altitude control group according to their clustering patterns and negligible Andean-specific genetic component, as pointed out by CHOMOPAINTER/fineSTRUCTURE and ADMIXUTURE estimates respectively of haplotypes sharing and individual ancestry proportions (Fig. 1c and Supplementary Results).

**Figure 2.**
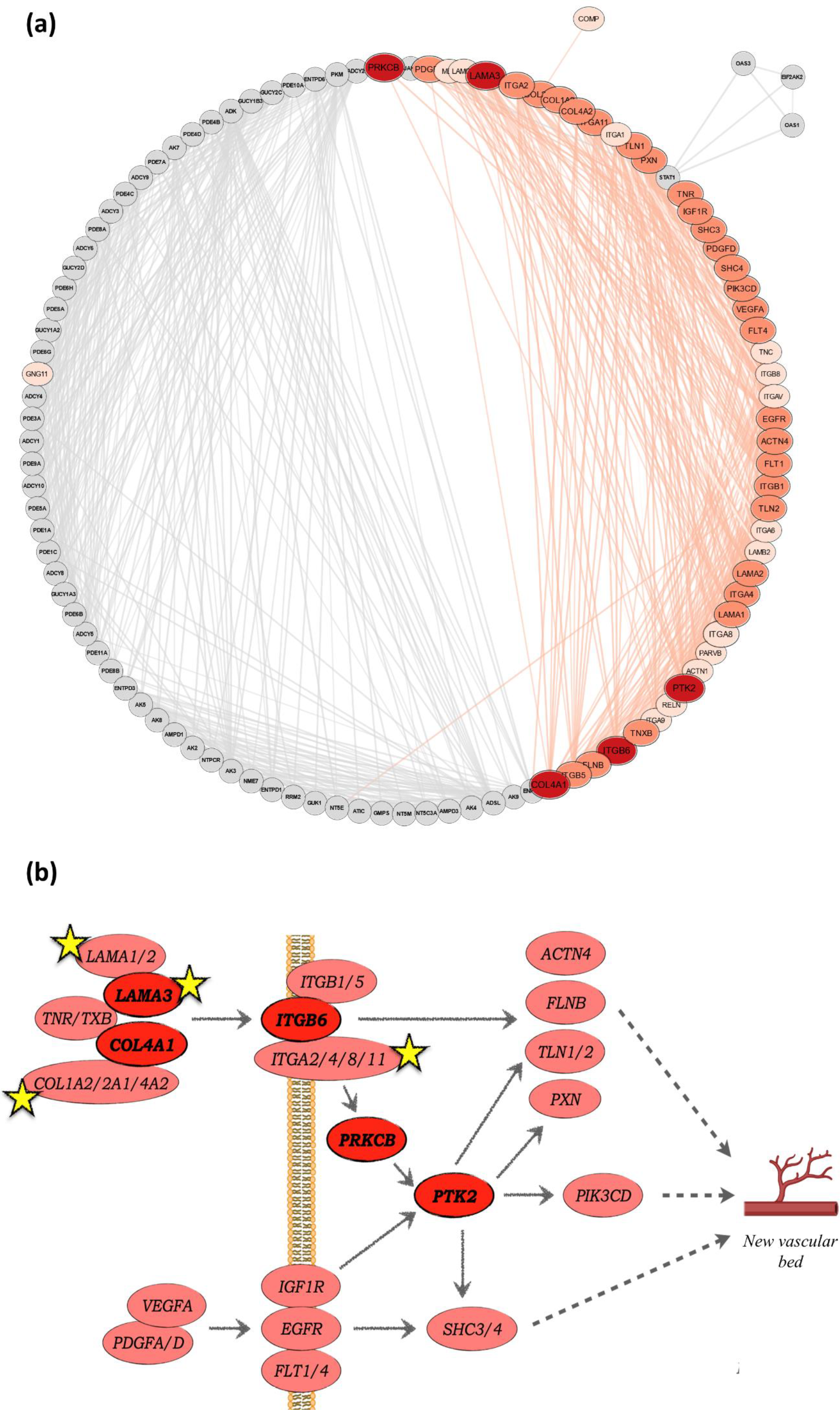
Functional relationships between genes putatively mediating polygenic adaptations evolved by Andean populations and their role in angiogenic processes. **(a)** Circular network build with Cytoscape v3.9.1 and displaying protein-protein functional interactions inferred for the entire set of adaptive genes belonging to significant pathways identified by *signet* analyses according to both WGS and imputed datasets. Gene products that establish similar connections in the network are placed next to each other, comporting the delineation of two well-distinguishable functional clusters. In detail, immune-related genes (e.g., belonging for example to the *Herpes simplex virus 1 infection* pathway) are reported as light-grey circles, while angiogenesis-related genes (i.e., belonging to the *Focal adhesion* pathway) are displayed with dark-red/dark-pink ellipses. Light-pink ellipses instead indicate those genes belonging to the *Focal adhesion* pathways for which involvement in the induction and regulation of vascular sprouting is not well established. **(b)**. Scheme displaying functional interactions between *Focal adhesion* adaptive genes contributing to improved placental/embryo angiogenesis in Andean populations. The most robust candidate adaptive loci, supported by both nSL-based and H12-based network analyses are reported as dark-red ellipses, while genes supported by a single statistic are displayed as pink ellipses. During placenta and embryo development, extracellular matrix components (e.g., *LAMA3*) interact with integrin-receptors (e.g., *ITGB6*) stimulating protein kinases (e.g., *PRKCB* and *PTK2*), which subsequentially activate those signalling cascades essential for migration of endothelial cells and formation of new vascular structures. The yellow stars mark those genes previously identified as loci putatively mediating polygenic adaptation to hypobaric hypoxia in Tibetan/Sherpa populations, thus providing evidence for partial genetic convergence between Andean and Himalayan adaptive traits in addition to the significant convergence observed at the biological function/pathway level.

### Genomic Signatures Ascribable to Polygenic Adaptive Events in Andean Populations

Genome-wide distributions of the number of segregating sites by length (nSL) and H12 statistics, which are informative of various types of selective events, were calculated for Andean and control groups relying on both WGS and imputed data. Finally, we used these distributions as input for the *signet* algorithm to infer gene networks made up of loci belonging to the same functional pathway and that have been concurrently targeted by natural selection (see Materials and Methods). By considering only pathways confirmed by each selection statistic and by both WGS and imputed datasets, we shortlisted the most plausible biological functions having evolved adaptively in the considered populations. Significant gene-networks identified for the low-altitude control group were found to participate primarily to immune-related pathways (e.g., *Staphylococcus aureus infection, Human papillomavirus infection*, and *Arachidonic acid metabolism*), suggesting the evolution of adaptations able to mediate immune and inflammatory responses to pathogens (Supplementary Table 2). Similar selective pressures seem to have acted also on Andean groups, as attested by their putative adaptive loci belonging to the *Purine metabolism* and *Herpes simplex virus 1 infection* pathways (Fig. 2a and Supplementary Table 3).

Conversely, adaptive events plausibly contributing to the response to stresses imposed by the high-altitude environment were inferred only in Andeans and were mediated by genes involved in the *Focal adhesion* pathway, which regulates cells adhesion to extracellular matrix and subsequent signal transduction at the intracellular level (Fig. 2a-b and Supplementary Table 3). Most of the loci constituting the significant networks identified by analyzing both Aymara WGS and Andean imputed datasets are indeed known to play a pivotal role in angiogenetic processes (Fig. 2 a-b). Especially five of these genes (i.e., *LAMA3, COL4A1, ITGB6, PRKCB*, and *PTK2*) showed patterns ascribable to polygenic adaptive evolution according to both nSL-based and H12-based network analyses, thus representing the most robust candidate targets of positive selection (Fig. 2b). Among them, *ITGB6, PRKCB*, and *PTK2* have been proved to be the main regulators of vascular sprouting, which implies the transformation of blood vessels endothelial cells into cells able to migrate and proliferate out of the vessel to create new vascular structures (Kanehisa & Goto, 2000; Jufri et al., 2015; Cao et al., 2019; Daniunaite et al., 2021; Sayers et al., 2022; Wang et al., 2022; You et al., 2022; Chen et al., 2023). In fact, *PTK2* inactivation has been demonstrated to prevent the beginning of the angiogenetic process, leading to a lack of development of vascular tissue in embryos and thus to their premature death (Shen et al., 2005; Adams & Alitalo, 2007). Although overall adaptive evolution of loci at the *Focal adhesion* pathway was highlighted by significant networks obtained according to both nSL and H12 statistics, the remaining 28 genes included in these networks were pointed out as potentially adaptive by a single statistic. Nevertheless, their functional relationships with the highly confirmed loci described above have been well established and they are known to contribute to the regulation of blood vessels formation as well (Fig. 2 a-b). For instance, multiple studies collected evidence supporting the interaction among *PTK2, VEGFA, PDGFA*, and *PDGFD* as a key driver of blood vessels formation in placenta, and some variants at these loci are found to be associated with reduced functionality of maternal-fetal circulation, ischemic placental disease, and impaired fetal growth (Khankin & Karumanchi, 2010; Russell et al., 2019)

### Convergent Adaptive Evolution Between Andean and Himalayan Populations

Interestingly, *LAMA1/2/3, COL1A2/2A1/4A2*, and *ITGA2/4/8/11* have been previously found to participate to gene networks targeted by natural selection in Tibetan and Sherpa populations^34^. Such an incomplete genetic convergence between Andean and Himalayan high-altitude peoples is not unexpected within the framework of an evolutionary scenario largely characterized by polygenic adaptive events, in which single loci play a negligible role with respect to their overall synergic interactions. Nevertheless, we provided evidence for the same biological functions/pathways (i.e., those related to angiogenetic processes) having adaptively evolved in both the considered populations, plausibly in response to the selective pressure imposed by hypobaric hypoxia. The identified genetic signatures might indeed contribute to the increase in the density of blood vessels that results in an improved blood flow and oxygen delivery to tissues despite the hypoxic stress, which is an adaptive trait qualitatively (but not quantitatively) similar between Andean and Himalayan groups (Gnecchi-Ruscone et al., 2018; Sharma et al., 2022; Wu et al., 2022). In fact, such a trait appears to be more enhanced and generalized in Tibetan/Sherpa populations with respect to Andean ones, being especially limited to improved uterine artery blood flow during pregnancy in the latter groups (Beall, 2007; Julian & Moore, 2019). The fact that several of the adaptive loci identified in the present study (e.g., *PTK2, VEGFA, PDGFA*, and *PDGFD*) play a pivotal role in the development of vascular tissue specifically in placenta and embryo, and thus in the establishment of efficient maternal-fetal circulation (Shen et al., 2005; Adams & Alitalo, 2007; Khankin & Karumanchi, 2010; Russell et al., 2019), seems to support such an observation. Moreover, it might suggest that natural selection succeeded in optimizing angiogenesis in Andean individuals mainly in early life and/or at the reproductive stage, thus leading to improved fetal development during intrauterine life (e.g., in terms of maturation of the respiratory system), which represents a key aspect to reduce neonatal mortality at high altitudes where efficiency of respiratory functions is crucial to ensure the individual’s survival.

In conclusion, the obtained results highlighted polygenic mechanisms mediated by adaptive evolution of several focal adhesion genes as the possible drivers of enhanced angiogenesis and oxygen transport in Andean human groups similarly to what previously observed in Tibetan/Sherpa populations, thus providing unprecedented evidence for the molecular bases of their convergent adaptation to hypobaric hypoxia.

## Materials and Methods

### Assembled Datasets

A WGS dataset including 9,439,267 SNVs for 24 Bolivian Aymara individuals was assembled from the data generated by (Lindo et al., 2018) and used to perform selection scans. Genome-wide genotype data were also collected for 141 individuals belonging to different Andean high-altitude populations to perform validation analyses. In detail, 713,014 genetic variants were retrieved for 26 Bolivian Aymaras, as well as for 21, 22 and nine subjects respectively from Aymara, Quechua and Uros populations settled in the Peruvian Titicaca Lake area (Gnecchi-Ruscone et al., 2019). Genotypes for 364,470 loci were collected also for 40 Quechua and 23 Aymara additional individuals (Reich et al., 2012).

### Data Curation and Imputation of the Validation Dataset

WGS data were submitted to QC procedures to filter for high-quality genotypes and to exclude possible closely related individuals. Specifically, the PLINK package v2.0 (Chang et al., 2015) was used to retain only SNVs on autosomal chromosomes, variants/samples showing missing data < 5%, and loci with non-significant deviations from the Hardy-Weinberg equilibrium (p > 1.059 x 10^−9^). We also removed variants showing ambiguous A/T or G/C alleles and we calculated pairwise identity-by-descent (IBD) statistics by estimating the genome-wide proportions of shared alleles for each pair of individuals. Then, to exclude subjects related to the second degree, we filtered out one individual from each pair showing an IBD coefficient > 0.270, as previously proposed for populations of Native American ancestry (Ojeda-Granados et al., 2022) (Supplementary Methods). This led to the generation of a high-quality WGS dataset including 7,966,198 SNVs and 24 individuals.

The described QC filters were applied also to the two assembled genome-wide datasets, making available for validation analyses 681,932 and 364,413 variants, respectively. These data were merged by selecting only the 210,540 variants shared between both datasets and were phased by means of the Eagle algorithm v2.4 (Loh et al., 2016) implemented in the Michigan Imputation Server (MIS) v1.2.4 (Das et al., 2016) using the 1000 Genomes Project Phase 3 and WGS datasets as reference panels. The MIS Minimac4 algorithm was then used to impute genotypes for 10,715,833 variants, which were submitted to the previously described QC procedures. In addition, the *bcftools* package v1.8 (Li et al., 2009) was used to retain only loci showing a squared Pearson correlation coefficient (r^2^) between imputed genotype probability and true genotype call > 0.95 (Palmer & Pe’er, 2016). Finally, IBD kinship coefficients were calculated for each pair of individuals in the imputed dataset using the same approach described for WGS data. According to these post-imputation QC steps, a high-density imputed dataset including 130 Andean individuals characterized for 8,024,216 variants was obtained.

### Assessing Samples’ Genetic Ancestry and Consistency Between Non-imputed/Imputed data

To verify that the collected Bolivian Aymara WGS were representative of the genetic variation observable across Andean high-altitude populations, we first explored their distribution within the overall genomic landscape of Native American populations. For this purpose, the high-quality WGS dataset was merged with a reference panel made up of genome-wide genotypes from un-admixed individuals from Central and South American indigenous populations (Gnecchi-Ruscone et al., 2019), and was submitted to Principal Component Analysis (PCA). Concurrently, to test also for consistency between non-imputed and imputed data, we performed PCA again by substituting the imputed data to the original ones as concerns the Andean populations included in the used Native American reference panel. In doing so, the *smartpca* method implemented in the EIGENSOFT package v6.0.1(Patterson et al., 2006) was used after having filtered the dataset for variants in high linkage disequilibrium (LD). LD pruning was obtained by considering sliding windows of 50 nucleotides sites in size and advancing by five loci at the time, as well as by removing variants for each pair showing r^2^ > 0.2. Moreover, IBS-based matrices of individual pairwise differences in the genome-wide proportion of shared alleles were computed on both the non-imputed and imputed LD-pruned Andean datasets. These matrices were then compared by means of the Mantel correlation test (Mantel, 1967) using functions implemented in the R *vegan* package and empirically assessing statistical significance by running > 10,000 permutations.

### Selecting Low-altitude Control Populations

To assess whether selection signatures detected in Andean people are informative about adaptations in response to high-altitude selective pressures, we selected low-altitude control populations of South American ancestry from the previously described reference panel of Native American groups (Gnecchi-Ruscone et al., 2019) to be used for replicating selection scans. In doing so, our rationale was to identify South American groups that: i) share an ancient common origin with Andeans, ii) do not show evidence of remarkable admixture with Andeans occurred after the divergence from their common ancestors, iii) do not have experienced high-altitude-related selective pressures during their evolutionary history. For this purpose, we calculated outgroup-*f3* statistic in the form of *f3*(X, Andean; CHB) to formally assess the extent of shared drift between Andean populations and the other Central and South American populations (X). This was done using the *qp3Pop* function implemented in the ADMIXTOOLS package v3.0 (Patterson et al., 2012) by including only groups with more than five individuals and first by considering Andean populations separately (i.e., Aymara, Bolivian Aymara, Titicaca Aymara, Titicaca Quechua, and Titicaca Uros, respectively), and then by considering them as a whole (i.e., ANDEAN group).

Moreover, to accurately select the individuals to be included in the control group, we further investigated the genetic relationships between Andeans and the putative control populations pointed out by outgroup-*f3* analyses. To this end, we inferred individual-based genetic relationships by reconstructing haplotype sharing patterns by means of the CHROMOPAINTER/fineSTRUCTURE clustering approach (Lawson et al., 2012) and by estimating ancestry proportions for each subject using the ADMIXTURE model-based clustering algorithm (Alexander et al., 2009) (Supplementary Methods).

### Inferring Polygenic Adaptive Events from WGS and Imputed Data

A combination of selection scans suitable to detect both strong and weak selective events was applied to WGS and imputed data. In detail, the nSL statistic was computed for each variant using *selscan* v.1.1.0b (Szpiech & Hernandez, 2014) and a 20,000 bp threshold for gap scale, 200,000 bp as the maximum gap length, and 4,500 consecutive loci as the maximum extension parameter (Ferrer-Admetlla et al., 2014). The H12 statistic was instead calculated with an ad hoc script using sliding windows of 200,000 bp and advancing by one variant at the time as previously suggested (Garud et al., 2015). Unstandardized values for each statistic were then normalized across multiple frequency bins by subtracting to each of them the bin average score and by dividing the resulting value by the related standard deviation.

In order to test a realistic approximation of a polygenic adaptation model, we retained as the representative scores of a given gene, the highest normalized nSL and H12 values computed among those obtained for all variants annotated on that gene. The resulting genome-wide distributions of selection scores were then submitted to gene-network analyses using the R *signet* algorithm (Gouy et al., 2017) and by setting 20,000 iterations to generate null distributions of the highest scoring subnetwork (HSS) for each gene-network of a given size to be compared with the obtained HSS. This provided a p-value for all the identified gene networks and those < 0.05 after controlling the false discovery rate (FDR) were used to point out combinations of multiple genes that participate to the same functional pathway (according to information from the KEGG database (Kanehisa & Goto, 2000)) and that have been targeted by natural selection. The most plausible biological functions having evolved adaptively in the considered population groups were finally identified as those supported by i) both nSL-based and H12-based significant networks, ii) replication of the results obtained from WGS data when imputed data were submitted to the same pipeline of analyses.

## Supporting information

Supplementary Methods and Results

## Acknowledgements

We acknowledge support from the Fondazione Cassa di Risparmio in Bologna through the project “Genetic adaptation and acclimatization to high altitude as experimental models to investigate the biological mechanisms that regulate physiological responses to hypoxia”, which was granted to M.S. (n. 2019.0552). We also would like to thank John Lindo for having kindly shared Aymara WGS data without which this work would have not been possible.

## Author Contributions

G. F., A. I., S. S., Data curation, Software, Formal analysis, Investigation, Writing - original draft; P. A., K. S., C. O-G. Software, Formal analysis; S. D. F., M. I., A. D., P. T. S., P. C., M. T., M. d. M., G. A. G-R., L. N., A. C., G. M., D. P., Data curation, Writing - review and editing; P. G., D. L., D. P., Resources; M. S., Conceptualization, Resources, Supervision, Funding acquisition, Methodology, Writing - review and editing.

## Data Availability

WGS data that support the findings described in the present study are available at the NCBI Sequence Read Archive (accession n. PRJNA470966)^13^. Genotype data used for validation analyses are instead available at https://figshare.com/articles/South_American_dataset_Gnecchi-Ruscone_et_al_2019_/7667174^21^.

## Code Availability

All the codes used in the present study are available as functions that were either implemented in published R packages or have been previously provided at https://github.com/paoloabondio/Ojeda-Granados_et_al_2021.

