## Supplementary Table S1 and S2 for "Convergent evolution of complex adaptive traits enabled human life at high altitudes"

**Supplementary Table S1.** Native American populations considered in the present study. In bold are represented the high-altitude groups included in the whole genome sequence (WGS) and validation datasets encompassing a total of 165 individuals used to perform selection scans.

| Location | Population | N | Ref |
|---|---|---|---|
| **Bolivia** | **Aymara_WGS** | **24** | Lindo et al. 2018 |
| **Bolivia** | **Bolivia_Aymara** | **26** | Gnecchi- Ruscone et al. 2019 |
| **Peru** | **Titicaca_Aymara** | **21** | Gnecchi- Ruscone et al. 2019 |
| **Peru** | **Titicaca_Quechua** | **22** | Gnecchi- Ruscone et al. 2019 |
| **Peru** | **Titicaca_Uros** | **9** | Gnecchi- Ruscone et al. 2019 |
| **Peru/Bolivia** | **Quechua** | **40** | Reich et al. 2012 |
| **Chile/Bolivia** | **Aymara** | **23** | Reich et al. 2012 |
| Chile | Yaghan | 4 | Reich et al. 2012 |
| Chile | Hulliche | 4 | Reich et al. 2012 |
| Peru | Huambisa | 8 | Gnecchi- Ruscone et al. 2019 |
| Peru | Cashibo | 10 | Gnecchi- Ruscone et al. 2019 |
| Peru | Shipibo | 17 | Gnecchi- Ruscone et al. 2019 |
| Peru | Ashaninka | 10 | Gnecchi- Ruscone et al. 2019 |
| Peru | Yanesha_HS | 23 | Gnecchi- Ruscone et al. 2019 |
| Peru | Yanesha_IS | 23 | Gnecchi- Ruscone et al. 2019 |
| Argentina | Wichi | 24 | Gnecchi- Ruscone et al. 2019 |
| Argentina | Wichi2 | 5 | Reich et al. 2012 |
| Argentina | Chane | 2 | Reich et al. 2012 |
| Paraguay/Argentina | Guarani | 6 | Reich et al. 2012 |
| Brazil | Surui | 24 | Reich et al. 2012 |
| Brazil | Karitiana | 13 | Reich et al. 2012 |
| Brazil | Jamamadi | 1 | Reich et al. 2012 |
| Brazil | Parakana | 1 | Reich et al. 2012 |
| Brazil | Arara | 1 | Reich et al. 2012 |
| Colombia | Ticuna | 6 | Reich et al. 2012 |
| Colombia | Piapoco | 7 | Reich et al. 2012 |
| Colombia | Guahibo | 6 | Reich et al. 2012 |
| Colombia | Wayuu | 11 | Reich et al. 2012 |
| Colombia | Waunana | 3 | Reich et al. 2012 |
| Colombia | Embera | 5 | Reich et al. 2012 |
| Colombia | Kogi | 4 | Reich et al. 2012 |
| CostaRica | Teribe | 3 | Reich et al. 2012 |
| CostaRica | Cabecar | 31 | Reich et al. 2012 |
| CostaRica | Bribri | 4 | Reich et al. 2012 |
| CostaRica | Maleku | 3 | Reich et al. 2012 |
| Guatemala | Kaqchikel | 13 | Reich et al. 2012 |
| Mexico | Tzotzil | 36 | Gnecchi- Ruscone et al. 2019 |
| Mexico | Mixe | 17 | Reich et al. 2012 |
| Mexico | Mixtec | 5 | Reich et al. 2012 |
| Mexico | Zapotec1 | 22 | Reich et al. 2012 |
| Mexico | Zapotec2 | 21 | Reich et al. 2012 |
| Mexico | Tepehuano | 25 | Reich et al. 2012 |
| Mexico | Pima | 33 | Reich et al. 2012 |
| Canada | Chipewyan | 15 | Reich et al. 2012 |

**Supplementary Table S2.** Gene-networks identified as significant for the low-altitude control group by signet analyses based on nSL/H12 statistics.

**H1-2**

| Dataset | Pathway | Pathway size | Subnetwork size | Subnetwork Score | P-value | Subnetwork genes |
|---|---|---|---|---|---|---|
| Low-altitude control dataset | Arachidonic acid metabolism | 45 | 3 | 5.246 | 0.001 | *CYP2C8 GPX5 GPX6* |
| Low-altitude control dataset | Glutathione metabolism | 57 | 3 | 8.441 | 0.003 | *GGT5 GPX5 GPX6* |
| Low-altitude control dataset | Staphylococcus aureus infection | 74 | 4 | 2.001 | 0.004 | *C1QB C2 C3 CFB* |

**nSL**

| Dataset | Pathway | Pathway size | Subnetwork size | Subnetwork Score | P-value | Subnetwork genes |
|---|---|---|---|---|---|---|
| Low-altitude control dataset | PI3K-Akt signaling | 305 | 14 | 6.574 | 0.002 | *CSF1R EFNA5 ERBB4 FGF1 FGF2 FGFR2 FLT1 FLT3 ANGPT1 IGF1R IRS1 NTRK2 BDNF FGF18* |
| Low-altitude control dataset | Rap1 signaling | 190 | 14 | 6.500 | 0.003 | *CSF1R EFNA5 FGF1 FGF2 FGFR2 FLT1 FGF20 ANGPT1 IGF1 IGF1R PDGFRB PDGFC TEK FGF18* |
| Low-altitude control dataset | Human papillomavirus infection | 328 | 12 | 6.353 | 0.003 | *ITGA11 FN1 COL6A3 TNC IBSP LAMA1 LAMA4 LAMC1 LAMC2 SPP1 THBS4 TNR* |
