## Supplement Table S3 for "Convergent evolution of complex adaptive traits enabled human life at high altitudes"

**Supplementary Table S3.** Gene-networks identified as significant by signet analyses based on nSL/H12 statistics and conducted on Andean WGS and imputed datasets.

**H1-2**

| Dataset | Pathway | Pathway size | Subnetwork size | Subnetwork Score | P-value | Subnetwork genes |
|---|---|---|---|---|---|---|
| Andean imputed dataset | Purine metabolism | 19635 | 37 | 8.0919 | 0.0052 | *ADCY1 ADCY2 ADCY3 ADCY5 ADCY9 AK7 ADK ADSL ADCY4 AK2 FHIT ENPP4 AK5 AMPD3 GUCY1A2 GUCY1A1 GUCY1B1 NME7 GUCY2D ATIC NT5E PDE11A NT5C3A PDE1A PDE1C PDE3A PDE8A PDE4D PDE7A PDE8A PKM NT5M PDE8B PDE5A GMPS ENTPD1 ENTPD3* |
| Andean imputed dataset | Herpes simplex virus 1 infection | 19635 | 54 | 9.9671 | 0.0012 | *ZNF559-ZNF177 ZNF211 ZNF786 ZFP90 ZNF560 ZNF563 ZNF709 ZNF721 ZNF585A ZNF285 ZNF544 ZIK1 ZNF841 ZFP82 ZNF619 ZNF621 ZNF181 ZNF517 JAK1 ZNF699 ZNF568 ZNF773 EIF2AK4 ZNF716 ZNF727 OAS1 OAS3 ZNF44 ZNF416 ZNF398 ZNF529 STAT1 ZNF7 ZNF19 ZNF23 ZNF91 ZNF124 ZNF136 ZNF141 ZNF177 ZNF180 ZNF184 ZNF112 ZNF343 ZNF669 ZNF671 ZNF30 ZNF551 ZNF616 ZNF461 ZNF561 EIF2AK3 ZNF254 ZNF432* |
| Andean imputed dataset | **Focal adhesion** | 19635 | 18 | 5.5131 | 0.0438 | ***COL1A2 COL2A1 COL4A1 LAMA1 ITGB1 ITGB5 ITGB6 LAMA3 LAMB2 PRKCB RELN PTK2 PXN TLN1 TNR TNXB ACTN4 TLN2*** |
| Aymara WGS | Herpes simplex virus 1 infection | 20096 | 31 | 6.8867 | 0.0115 | *ZNF268 ZNF641 ZNF480 ZNF791 ZNF169 ZNF721 ZNF25 ZNF311 ZNF780A ZNF615 ZNF619 ZNF454 ZFP57 EIF2AK4 OAS1 OAS3 ZNF334 EIF2AK2 ZNF248 ZNF490 ZNF350 STAT1 ZNF3 ZNF10 ZNF43 ZNF141 ZNF669 ZNF614 ZNF300 ZNF254 ZNF432* |
| Aymara WGS | **Focal adhesion** | 20096 | 11 | 6.2476 | 0.0276 | ***COL4A1 COMP ITGA2 ITGA4 ITGB6 ITGB8 LAMA2 LAMA3 LIG4 PTK2 ITGA8*** |
| Aymara WGS | Purine metabolism | 20096 | 26 | 8.5079 | 0.0064 | *PDE10A ADCY3 ADCY5 AK8 ADCY4 AK9 FHIT ENPP4 AK5 AMPD1 GUCY1A2 GUCY2C GUK1 AK3 PDE11A PDE1A PDE1C PDE4D PDE6A PDE6G PDE6B PDE8B PDE5A GMPS ENTPD6 ENTPD3* |

| **nSL** | | | | | | |
|---|---|---|---|---|---|---|
| **Dataset** | **Pathway** | **Pathway size** | **Subnetwork size** | **Subnetwork Score** | **P-value** | **Subnetwork genes** |
| Andean imputed dataset | Purine metabolism | 14711 | 31 | 7.1848 | 0.0221 | *ADA ADCY2 PDE10A ADCY5 ADCY9 AK7 ADK AK8 ADCY4 AK4 FHIT AK5 AMPD3 GUCY1A2 GUCY1A1 NME7 GUCY2D ATIC AK3 NT5C3A PDE1A PDE4B PDE4D PDE6H PDE7A PDE9A ENPP1 RRM2 PDE8B PDE5A GMPS* |
| Andean imputed dataset | Herpes simplex virus 1 infection | 14711 | 26 | 6.7918 | 0.0316 | *ZNF813 ZNF420 ZNF558 ZNF569 ZNF383 ZNF169 ZNF585A EIF2AK1 ZNF850 JAK1 ZNF699 ZNF568 EIF2AK4 ZNF331 STAT1 ZNF3 ZNF665 ZNF528 ZNF566 ZNF616 ZNF468 ZNF765 ZNF845 ZNF300 ZNF585B ZNF561* |
| Andean imputed dataset | **Focal adhesion** | 14711 | 23 | 6.5541 | 0.0405 | ***LAMC3 COL4A1 COL4A2 ITGA11 FLNB PARVB TNC ITGA6 ITGA1 ITGA2 ITGA9 ITGAV ITGB6 ITGB8 LAMA2 LAMA3 LAMB2 PRKCB PTK2 PXN TNR ITGA8 ACTN1*** |
| Aymara WGS | **Focal adhesion** | 19392 | 12 | 5.6383 | 0.0236 | ***EGFR FLT1 FLTA4 IGF1R GNG11 SHC4 MET PDGFA PIK3CD SHC3 VEGFA PDGFD*** |
| Aymara WGS | Purine metabolism | 19392 | 22 | 7.4699 | 0.0048 | *ADCY10 ADCY2 ADCY3 ADCY4 ADCY6 ADCY8 ADCY9 AK3 AK5 AMPD3 FHIT GMPS GUCY1A2 NT5C3A NTPCR PDE1C PDE3A PDE4B PDE4C PDE4D PDE8A PDE8B* |
| Aymara WGS | Herpes simplex virus 1 infection | 19392 | 37 | 7.0004 | 0.0063 | *EIF2AK4 JAK1 STAT1 ZFP30 ZIM3 ZNF100 ZNF114 ZNF12 ZNF160 ZNF180 ZNF254 ZNF300 ZNF304 ZNF324 ZNF347 ZNF354A ZNF415 ZNF443 ZNF528 ZNF534 ZNF540 ZNF551 ZNF554 ZNF566 ZNF571 ZNF582 ZNF583 ZNF615 ZNF623 ZNF727 ZNF77 ZNF772 ZNF783 ZNF790 ZNF813 ZNF845 ZNF850* |
