## Supplementary Methods and Results for "Convergent evolution of complex adaptive traits enabled human life at high altitudes"

### Supplementary Results

#### Testing representativeness of the assembled datasets

To verify the representativeness of the considered samples and to explore their relationship patterns within the genomic landscape of Native American populations, we compared the assembled high-quality WGS and imputed Andean datasets to the genomic variation observed in a wide reference panel made up of genome-wide genotypes for several Native American populations from Central and South America[1] (Supplementary Table 1).

Results of Principal Components Analysis (PCA) showed that along the latitudinal cline of Native American genetic diversity described by PC2, the high-altitude Andean samples formed a tight population cluster remarkably distinct from lowland groups, even with respect to those from Peru and the Amazon basin, who are the most geographically close to them (Fig. 1a). In addition, PC1 pointed to some groups from Costa Rica and Brazil as genetic outliers when compared with the bulk of the considered samples (Fig. 1a).

#### Selecting low-altitude control populations

Outgroup-f3 statistics and clustering analyses based on both the unsupervised model-based approach implemented in the ADMIXTURE algorithm and the haplotype-based CHROMOPAINTER and fineSTRUCTURE methods were applied to the assembled reference dataset to further investigate genetic relationships between Andeans and other Native American groups. This approach was aimed at selecting a low-altitude control group of South American ancestry that share an ancient common origin with Andeans without evidence of remarkable admixture after the divergence from their common ancestor, and that do not have experienced high-altitude-related selective pressures during its evolutionary history.

Consistently with sharing a common genetic background, the results of outgroup f3-statistics in the form of *f3(X,Andean;CHB)* confirmed that each Andean population (i.e., Aymara, Bolivian Aymara, Titicaca Aymara, Titicaca Quechua, and Titicaca Uros) tend to share the highest amount of genetic relatedness with the other Andean groups (Supplementary Fig. 2). This is followed, by a progressive decline in the amount of shared genetic drift experienced, where the Amazonian populations from Peru (i.e., Cashibo, Shipibo, Huambisa, Yanesha, and Ashaninka) appeared as the next most affine to Andeans (Fig. 1b, Supplementary Fig. 2). In fact, in a complex scenario that possibly involved distinct migrations into South America, with population replacements and/or multiple contacts between Central and South American groups to account for the genetic structure observed East and West of the Andes, the Peruvian Amazonian populations have been modeled as belonging to a non-Andean lineage that diverged from high-altitude Andeans[1]. Despite confirming the role of the Andes as a sharp genetic barrier, some proportions of gene flow from high-altitude Andeans into lowland Peruvian Amazonian groups were previously suggested for certain groups. However, the overall extent of "Andean" admixture occurred in Peruvian Amazonians after the divergence from their common ancestors was estimated to be relatively low[1].

Consistently with that and with harboring a non-Andean Peruvian Amazonian specific ancestry component, when looking at the individual-based relationship patterns emerged from the haplotype-based fineSTRUCTURE clustering approach, almost all the Amazonian ethnic groups from Peru were indeed found to belong to the same genetic cluster (Fig. 1c). The sole exception was represented by Huambisa, which instead clustered more closely to groups from Brazil and Colombia. Furthermore, the Andean-specific ancestry component (displayed in blue in Fig 1c) was observed at appreciable levels only in the Yanesha (12.9%) and Shipibo (9.7%) ethnic groups, while was nearly absent in the remaining Peruvian Amazonian populations, such as Ashaninka (1.1%) and Cashibo (0.001%). According to this pattern of population genetic structure, both the Huambisa and Yanesha populations, along with those Shipibo individuals showing higher levels of Andean admixture, were excluded from the identified control group, which finally resulted in 24 Cashibo, Ashaninka, and Shipibo subjects belonging to the Peruvian Amazonian genetic cluster.

### Supplementary Methods

#### Data curation of the validation dataset

To exclude subjects related to the second degree, we filtered out one individual from each pair showing an IBD coefficient > 0.270, as previously proposed for populations of Native American ancestry[2], which experienced severe founder events and long-term isolation, being thus characterized by low effective population size and remarkable inbreeding levels[1].

#### Estimating haplotype sharing patterns and individual ancestry proportions

The merged dataset including Andeans, Central and South American groups was phased with SHAPEIT2 v2.r790[3] by applying default parameters and HapMap phase 3 recombination maps. The CHROMOPAINTERv2/fineSTRUCTURE pipeline was then applied by first estimating the mutation/emission and the switch rate parameters with ten steps of the Expectation–Maximization (EM) algorithm on a subset of chromosomes {4, 10, 15, 22} using every individual both as "donor" and "recipient." Then, we averaged the obtained values across chromosomes (weighting by the number of markers) and individuals, and we used the estimated mutation/emission and switch rate parameters to run CHROMOPAINTER again on all chromosomes, considering a parameter k = 50 to specify the number of expected chunks to define a region. This value was suggested to be preferable compared with the default value of k = 100 when considering closely related populations[4]. The obtained matrix of haplotype sharing "chunk" counts was summed up across all the 22 autosomes and submitted to the fineSTRUCTURE clustering algorithm version fs2.1[5]. In detail, we ran the fineSTRUCTURE pipeline by setting 1,000,000 "burn-in" MCMC iterations, followed by additional 2,000,000 iterations and sampling the inferred clustering patterns every 10,000 runs. Finally, we set 1,000,000 additional hill-climbing steps to improve posterior probability and merge clusters in a stepwise fashion. Individuals were hierarchically assembled into clusters until reaching the final configuration dendrogram.

The unsupervised model-based clustering approach implemented in ADMIXTURE v.1.22 was also applied on the same merged dataset including 43 Amerindian populations after LD-pruning, by testing from K = 2 through K = 10 ancestral components. For each K tested, we performed 50 independent ADMIXTURE runs with a different random seed to monitor convergence and only those with the highest log-likelihood were considered. Concurrently, we calculated cross-validation (CV) errors for each K to identify the most reliable number of genetic clusters fitting with the data, which resulted in K = 8.

### Supplementary Figures

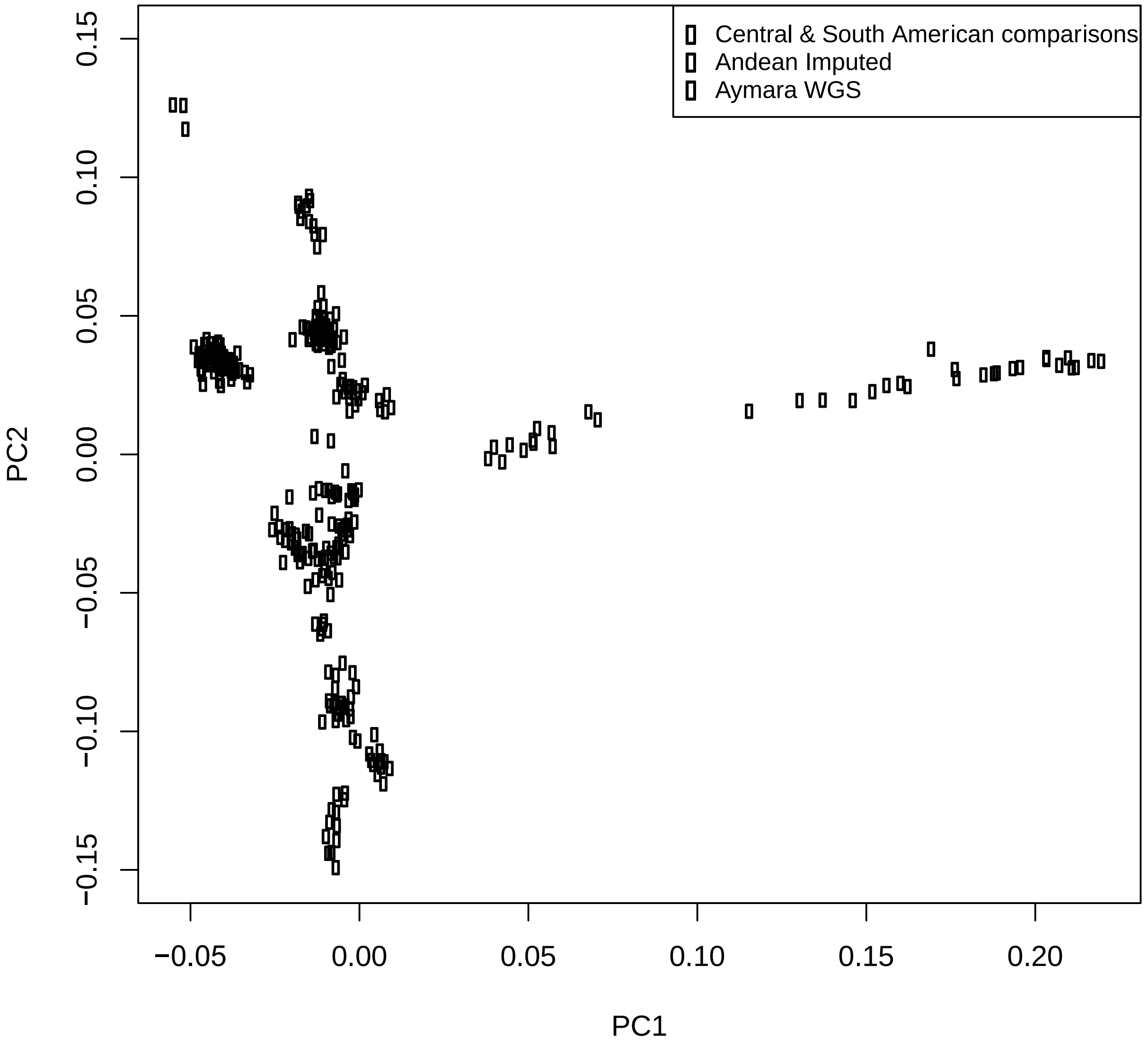


**Supplementary Figure 1.** PCA comparing Aymara WGS and Andean imputed datasets with the overall genomic landscape of Native American populations from Central and South America. Aymara individuals sequenced for the whole genome and Andean subjects characterized by imputed genome-wide data are represented by full green and red circles respectively, while gray dots refer to the considered populations from Central and South America used for the sake of comparison and detailed in Figure 1.

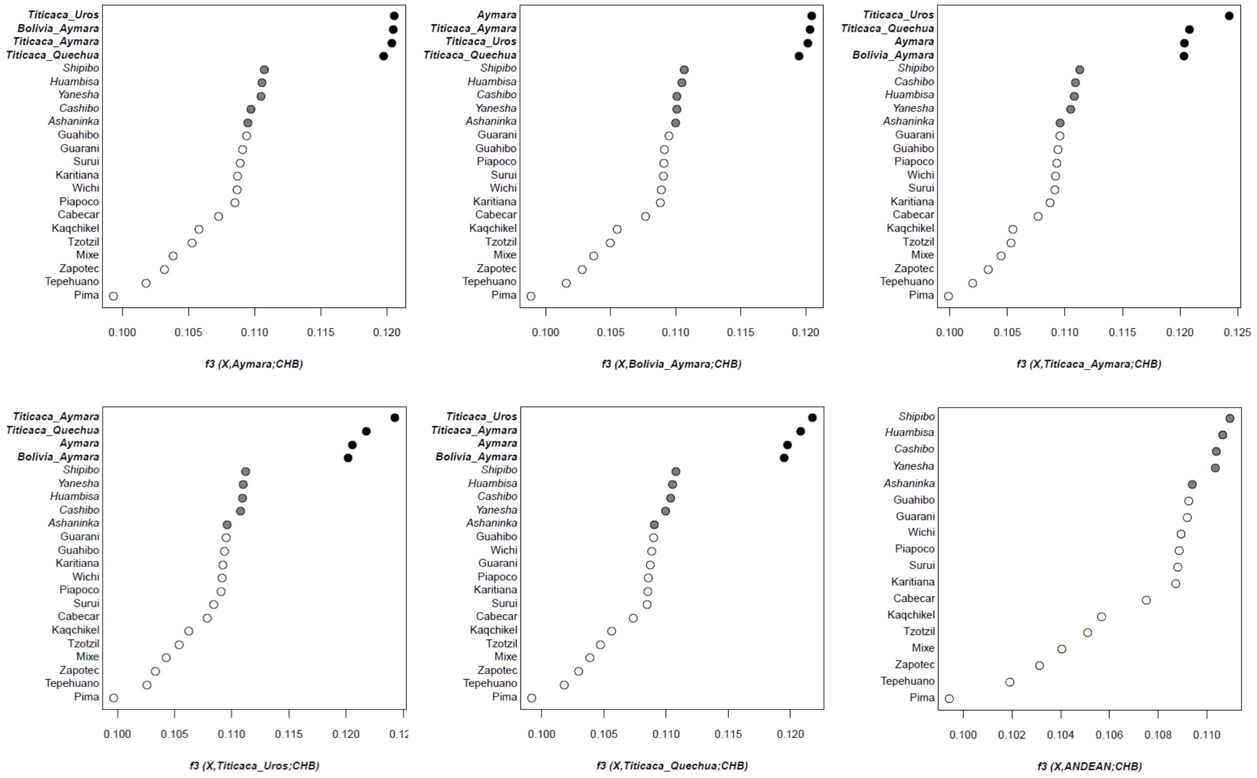


**Supplementary Figure 2.** Ranked values of the computed outgroup-f3 statistics showing patterns of genetic affinity between each Andean population and the other Central and South American groups included in the assembled reference dataset. In each plot, the other Andean populations compared with the test one, as well as the identified next most affine Amazonian populations from Peru, are represented by black and gray dots respectively.
